# SARS-CoV-2 Helicase might interfere with cellular nonsense-mediated RNA decay, insights from a bioinformatics study

**DOI:** 10.1101/2022.05.30.494036

**Authors:** Behnia Akbari, Ehsan Ahmadi, Mina Roshan Zamir, Mina Sadeghi Shaker, Farshid Noorbakhsh

## Abstract

Unraveling molecular interactions between viral proteins and host cells is key to understanding the pathogenesis of viral diseases. We hypothesized that potential sequence and structural similarities between SARS-CoV2 proteins and proteins of infected cells might influence host cell biology and antiviral defense. Comparing the proteins of SARS-CoV-2 with human and mammalian proteins revealed sequence and structural similarities between viral helicase with human UPF1. The latter is a protein that is involved in nonsense mediated RNA decay (NMD), an mRNA surveillance pathway which also acts as a cellular defense mechanism against viruses. Protein sequence similarities were also observed between viral nsp3 and human Poly ADP-ribose polymerase (PARP) family of proteins. Gene set enrichment analysis on transcriptomic data derived from SARS-CoV-2 positive samples illustrated the enrichment of genes belonging to the NMD pathway compared with control samples. Moreover, comparing transcriptomic data from SARS-CoV2-infected samples with transcriptomic data derived from UPF1 knockout cells demonstrated a significant overlap between datasets. These findings suggest that helicase/UPF1 sequence and structural similarity might have the ability to interfere with the NMD pathway with pathogenic and immunological implications.

## Introduction

SARS-CoV-2 proteome consists of structural and non-structural proteins (nsp) that play various roles in viral life cycle, from cell entry to viral replication and gene expression (1, 2). In addition to harboring elements necessary for their life cycle, viruses are known to possess the ability to interfere with host defense mechanisms. These mechanisms are quite diverse, with some viruses interfering in immunological defense mounted by leukocytes, while others inhibiting innate factors which restrict viral replication inside target cells. Producing proteins which bear sequence or structural similarities with host proteins is one of the mechanisms that viruses harness in order to influence anti-viral defense mechanisms. In the case of DNA viruses, which contain relatively big genomes, these virus-host protein similarities are sometimes the consequence of ‘gene hijacking’ which has occurred during viral evolution (3). For viruses with smaller genomes, and hence lower capacity to incorporate entirely new genes, other mechanisms might enable viral proteins to mimic host proteins (4). In this study we sought to identify potential similarities between SARS-COV-2 proteins with human proteins and investigate whether these similarities might potentially influence virus-host interactions. To this end, we first compared all SARS-COV-2 protein sequences with human reference protein sequences. This was followed by examining the structural similarities between some of these proteins. Next, using transcriptomic data derived from SARS-COV-2-infected cells/tissues we sought to investigate whether the observed similarities might have had an effect on host anti-viral mechanisms.

## Methods

### Data extraction

Reference protein sequences of SARS-CoV-2 were extracted from NCBI protein database (https://www.ncbi.nlm.nih.gov/protein/). Polyprotein and redundant sequences were removed. All accession numbers and their official gene symbols were shown in Table S1. 3D structure of proteins (2IYK, 2XZO, 2XZP, 6JYT, 5WWP) were downloaded from Protein Data Bank (PDB: https://www.rcsb.org/). RNA sequencing datasets were obtained from NCBI Gene Expression Omnibus (GEO: https://www.ncbi.nlm.nih.gov/gds). Datasets in which cells or patients were under treatment were excluded. The raw count data were downloaded in .csv or .txt format.

### Sequence and structural alignments

Protein sequence similarity searches were performed for all SARS-CoV-2 proteins against human and mammalian reference protein sequences using NCBI DELTA-BLAST (Domain Enhanced Lookup Time Accelerated BLAST: https://blast.ncbi.nlm.nih.gov/Blast.cgi). All DELTA-BLASTs were followed by three iterations of PSI-BLAST (Position-Specific Iterative-BLAST). DELTA-BLAST parameters are shown in Table S2. To report the strongest similarities, highest E-value/alignment score pairs were extracted for each query. Results were visualized as negative Log10 of E-values and alignment score. Structural alignment for helicase protein of SARS-CoV-2 against UPF1 and CH domain of UPF1 was carried out using Dali server (http://ekhidna2.biocenter.helsinki.fi/dali/).

### PPI network

The search tool for retrieval of interacting genes (STRING) (https://string-db.org/, version 11.0) database were applied to predict functional interactions of proteins with UPF1. Active interaction sources, experiments, and databases, as well as species limited to “Homo sapiens” and an interaction score□>□0.9 were applied to construct the PPI networks. Cytoscape software (version 3.6.1) was used to visualize the PPI network.

### Gene Ontology analysis

Functional enrichment analysis was performed using the Gene Ontology (GO) database (http://geneontology.org) and ClusterProfiler R package. GO term enrichment was done for biological processes (BP), molecular function (MF) and cellular components (CC) categories. Data were visualized using R ggplot2 package. Terms with adjusted p-value less than 0.05 reported significant.

### RNA-seq data processing

Datasets including RNA seq data from SARS-CoV2-infected samples were extracted from NCBI GEO. Datasets were imported into R studio using built-in “read.csv” function. Digital gene expression lists were generated using edgeR package and “DEGList” function. Data filtration and normalization were performed using Trimmed Mean of M-values (TMM) method via “calcNormFactors” function in edgeR package. Differentially expressed genes (DEGs) were determined from each dataset. T-test were performed to assess differential expression of genes between COVID-19 samples and healthy controls.Benjamini Hochberg method were used for p-value adjustment. To determine the effect size for each gene, the mean ratio of each COVID-19 gene versus the average expression of it in healthy controls were calculated (Fold-Change). Genes with adjusted p-value<0.05 and |log2 FC| > 1 were considered as DEGs. Furthermore, samples were annotated with NCBI official gene symbols using Homo sapiens annotation package (hgu133plus2.db). The biomaRt package were further used to match Ensembl gene IDs to official gene names extracted from hgu133plus2.db.

### Gene Set Enrichment Analysis

DEGs were extracted from datasets according to the above-mentioned pipeline. In the next step, DEGs from each dataset were sorted according to their FC. GSEA analyses were performed using the “GSEA” function of clusterProfiler package in R software. C2 category of the Msigdb was used as gene sets in our GSEA. p-values less than 0.05 were considered to indicate significant enrichment. GSEA results were visualized with “gseaplot2” function of R software’s enrichplot package.

## Results

### SARS-CoV-2 proteins show sequence similarities with human and mammalian proteins

In order to find potential protein sequence similarities between SARS-CoV-2 and Homo sapiens/ mammalian proteins, SARS-CoV-2 reference protein sequences (RefSeq) were extracted from NCBI protein database (S1Table). DELTA-BLAST (Domain Enhanced Lookup Time Accelerated Basic Local Alignment Search Tool) was used to compare all SARS-CoV-2 RefSeq protein sequences against reference sequences of Homo sapiens and mammalian proteins. As shown in Figure 1, SARS-CoV-2 non-structural protein 3 (nsp3) showed significant similarities (based on e value) with both human and mammalian proteins (matched with MACROD2 in iteration 1 and PARP15 in iteration 3). Interestingly, several groups have previously reported that nsp3 of coronaviruses interferes with interferon signaling pathway, possibly via the reversal of protein ADP-ribosylation, a posttranslational modification catalyzed by host poly (ADP-ribose) polymerases (PARPs) (5). Another significant result was detected for SARS-COV-2 helicase protein (non-structural protein 13) when aligned with either human or mammalian Refseq proteins. Viral helicase showed the highest similarities with UPF1 and Mov10L1 in humans and DNA2 in mammals, proteins which have helicase activity. Less significant similarities were observed between viral nucleocapsid phosphoprotein (with human MACF1 and Dystonin isoform 1eA precursor and mammalian SRRM5), viral surface glycoprotein (with human PSMA2), and viral nsp6 (with ADGRL3 in humans and MRM2 in mammals) and human/mammalian proteins (Fig 1A-D). Focusing on nsp3 and helicase, we next looked at all human proteins which show sequence similarities with these two proteins. Viral helicase (YP_009725308.1) exhibited higher degrees of sequence similarity with human UPF1 (UPF1 RNA helicase and ATPase), DNA2 (DNA replication ATP-dependent helicase/nuclease), MOV10L1 (Mov10 like RISC complex RNA helicase 1) and SMUBP2 (DNA-binding protein SMUBP-2), all members of helicase superfamily (Figs 2A and 2B) (6). Of these proteins, UPF1 has known roles in non-sense mediate RNA decay (NMD), a surveillance mechanism for mRNAs containing premature termination codons, as well as viral RNAs (Supp figure 1) (7). For viral nsp3, human MACROD1, MACROD2 as well as PARP9/15 (poly ADP-ribose polymerase family members) revealed the highest degrees of similarities. These proteins are mainly known as members of macro domain protein family which can bind to ADP-ribose (8) (Figs 2C and 2D).

**Figure 1.**
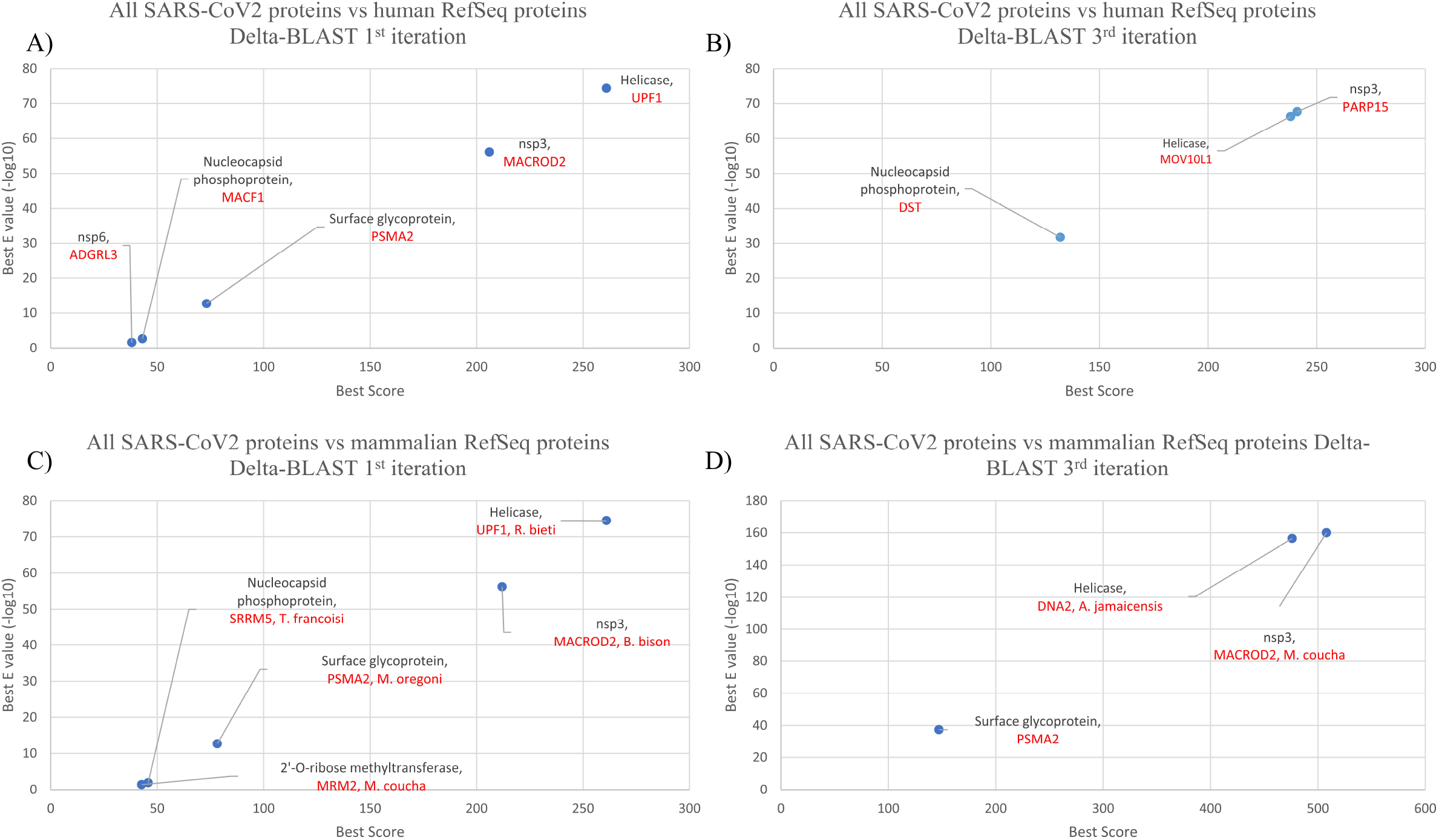
SARS-CoV-2 Nsp3 and helicase reveal higher sequence similarities with human/mammalian proteins. Results of the 1^st^ (A) and 3^rd^ (B) iterations of DELTA-BLAST comparing SARS-CoV-2 protein sequences with Homo Sapiens RefSeq proteins have been shown as negative log E values/alignment scores. Likewise, results of the 1^st^ (C) and 3^rd^ (D) iterations of DELTA-BLAST comparing SARS-CoV-2 proteins against mammalian RefSeq proteins are shown. Below every SARS-COV-2 protein, the name of the aligned human or mammalian protein (together with the species for the latter) is shown with red fonts. For each query only the highest log E value/score has been shown.

**Figure 2.**
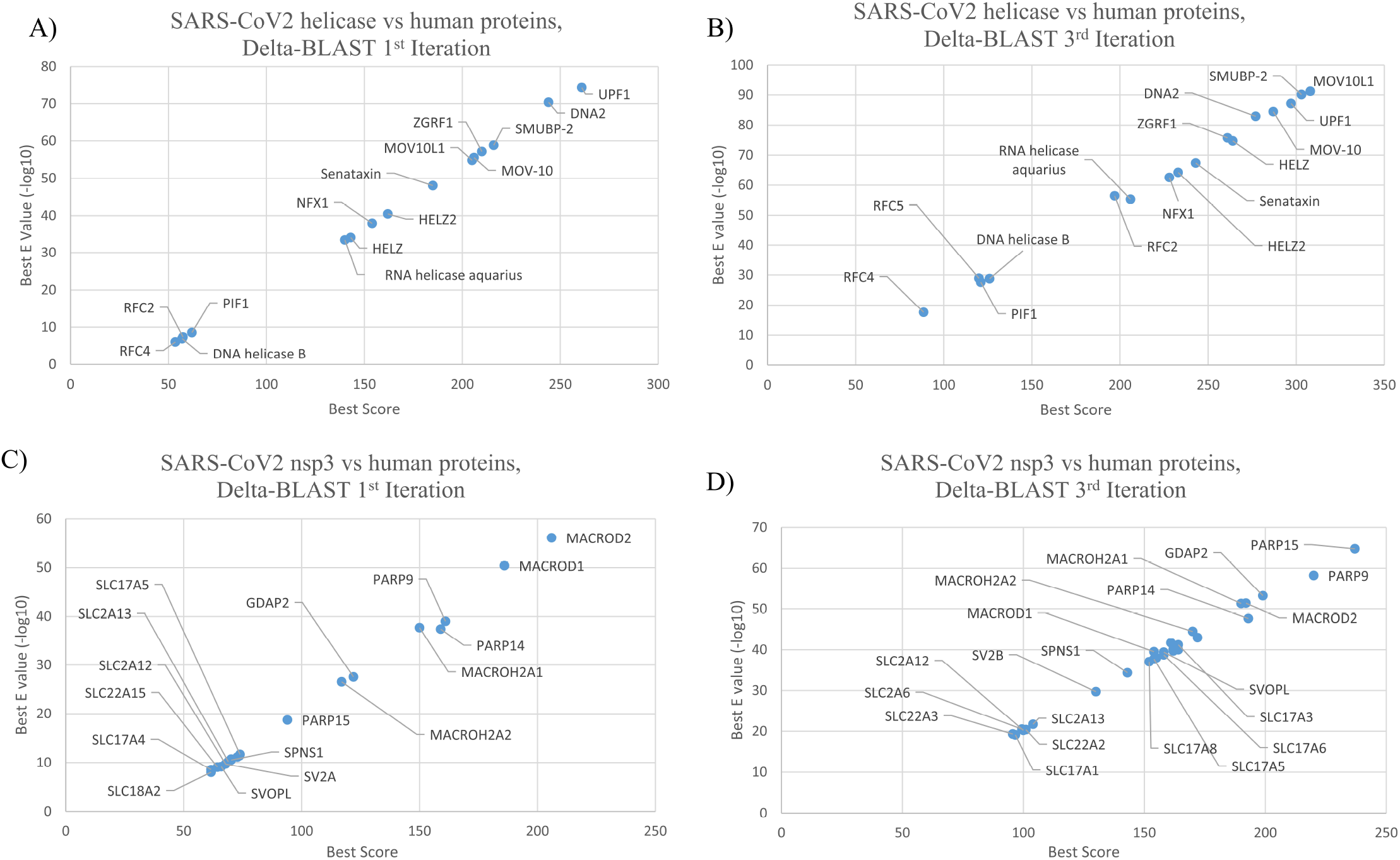
SARS-CoV-2 helicase protein sequence similarities with human proteins. Results of the 1^st^ and 3^rd^ iterations of DELTA-BLAST comparing SARS-CoV-2 helicase (A and B) or nsp3 (C and D) with Homo Sapiens RefSeq proteins.

### Helicase domain and CH domains of viral nsp13 show structural similarities with UPF1

To explore whether observed sequence similarity in SARS-CoV-2 helicase with human proteins is also associated with structural similarities, we compared SARS-CoV-2 helicase structure determined by Newman et al (PDB/6ZSL) with human UPF1 structure (PDB/2XZP) using DALI server (9). This structural comparison yielded a Z-Score of 26.6, and RMSD value of 3.4 (Fig 3A). We also compared helicase structure from two other highly pathogenic coronaviruses, i.e. SARS-COV and MERS-COV with human UPF1. The results showed high Z-score/RMSD values for these two viral helicases as well, indicating that the observed similarity was not limited to SARS-CoV-2 helicase (Fig 3A). UPF1 has 3 main domains: CH, Helicase and SQ domains (Fig 3B). CH domain interacts with UPF2 while SQ domain exerts an inhibitory effect on helicase. However, viral helicases include only helicase and CH domains and lack the inhibitory SQ domain (Fig 3B) (10). CH domain inhibits UPF1 catalytic activity, and this inhibition is relieved after binding of CH domain to UPF2, another NMD pathway member. Hence UPF1/UPF2 interaction via UPF1’s CH domain is a critical step in initiation of NMD pathway. To find out whether UPF1 CH domain has any structural similarities corresponding domain in viral helicase, we compared the crystal structure of this domain (PDB/2IYK) with viral helicase structures using DLAI server. As shown in Fig 3A, all three viral helicases showed a relatively high degree of similarity with UPF1 CH domain.

**Figure 3.**
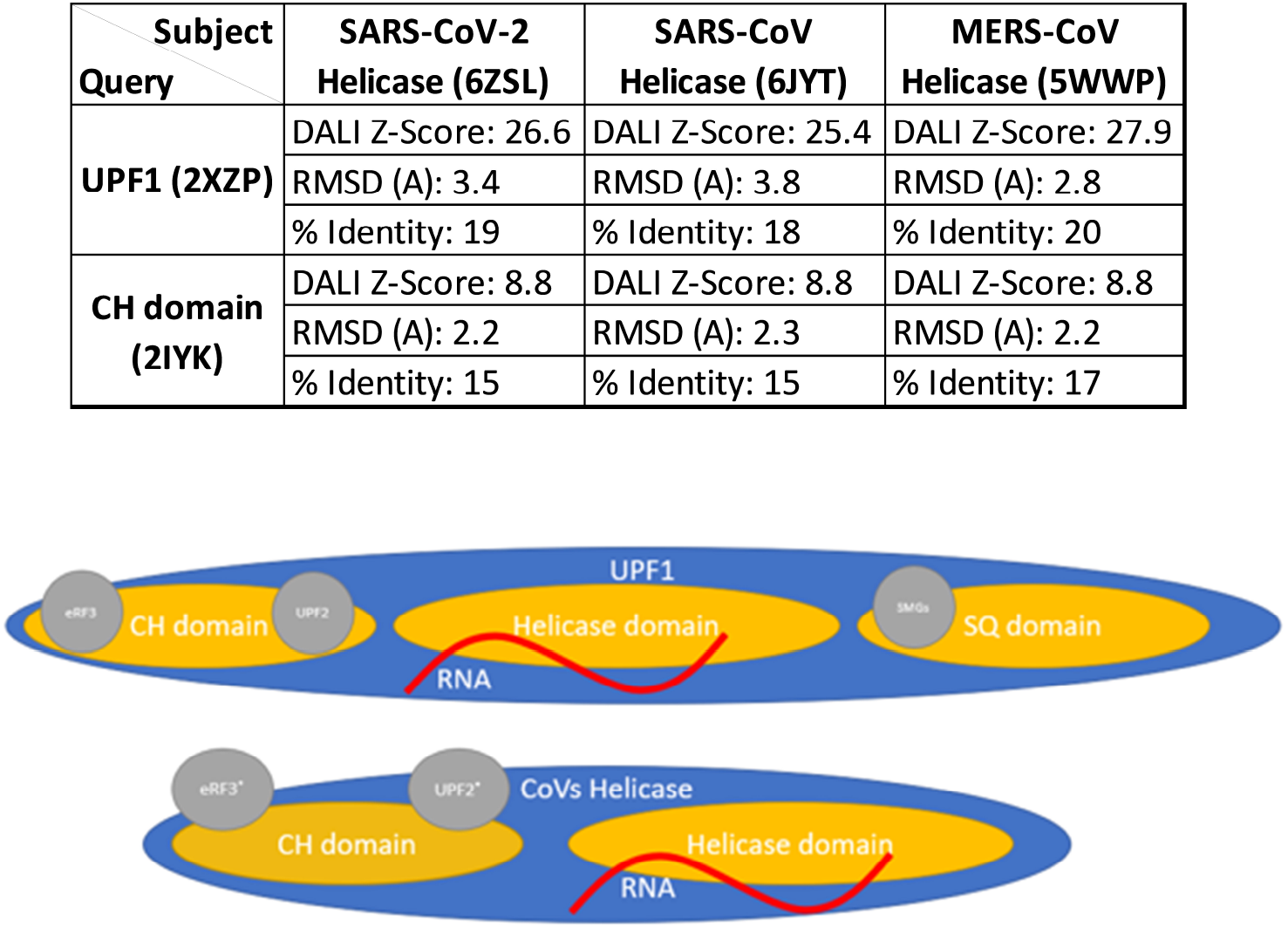
Structural comparison and schematic representation of domains of human UPF1 and coronaviral helicases. Results derived from DALI-server pairwise 3D domain comparison of SARS-CoV-2, SARS-CoV, and MERS-CoV helicase protein with UPF1 (A). Helicase core of viral helicase protein has a better score than the CH domain in comparison to UPF1. rmsd: root-mean-square deviation. Schematic representation of domains of human UPF1 and coronaviral helicases (B). UPF1 contains three domains; CH, helicase, and SQ domains, whereas coronaviral helicases lacks SQ domain.

### SARS-COV-2 infection leads to altered expression of NMD pathway transcripts

Nonsense-mediated decay (NMD) is a highly conserved pathway which exists in all eukaryotic organisms. NMD’s key function is to remove mRNAs which have been transcribed from genes with nonsense mutations, hence preventing these mRNAs from producing incomplete/truncated proteins (11).This is performed by detection of premature termination codons (PTCs) in mRNA transcripts (12). That said, NMD has been shown to target other ‘unwanted’ RNAs as well, including viral RNAs (13). While viral RNAs do not contain PTCs *per se*, the multi-cistronic nature of viral genomes makes their transcripts susceptible to NMD, as the internal stop codons are recognized as PTCs by the NMD pathway (7). Interestingly, different viruses have developed mechanisms to circumvent the targeting of their RNAs by NMD (14, 15). Considering the protein sequence/structure similarities that we observed between SARS-COV-2 helicase and human UPF1, we asked whether this might lead to any interference with host cells NMD pathway. To check whether NMD pathway members are affected in SARS-COV-2-infected cells/tissues, we decided to analyze RNA seq data derived from cells/tissues infected with the virus. We extracted raw data from for datasets deposited in NCBI GEO, i.e. GSE155974, GSE171110, GSE157103 and GSE182917. Of these. GSE155974 contains transcriptomic data from in vitro infected cells. GSE171110 and GSE157103 are derived from RNA seq analysis of cells obtained from COVID-19 and control patients and GSE182917 is derived from lung autopsy tissue (details are shown in Table S3). To investigate whether viral infection has influenced NMD pathway, we performed Gene Set Enrichment Analysis (GSEA) on these datasets, as described in the Methods section.

Our analyses showed negative enrichment of NMD pathway gene set members in the first three datasets; i.e. infected nasal organoid cells (Fig 4A), whole blood (Fig 4B) and leukocytes (Fig 4C) derived from COVID19 patients. In contrast, the last dataset; i.e. lung autopsy tissues, showed a positive enrichment for NMD gene set (Fig 4D). While airway epithelial cells are known as the major targets of SARS-CoV-2, monocytes and macrophages have also been shown to be infected by the virus, likely in a non-replicative manner (16, 17). NMD pathway members are known to restrict viral replication (18) and downregulation of NMD pathway elements has been shown to be associated with higher levels of viral proteins in infected cells (12). We believe that these data show potential suppression of NMD pathway activity in SARS-CoV-2-infected cells. Nonetheless, these transcriptional changes do not necessarily reflect perturbed UPF1 function caused by viral proteins; e.g. nsp13.

**Figure 4.**
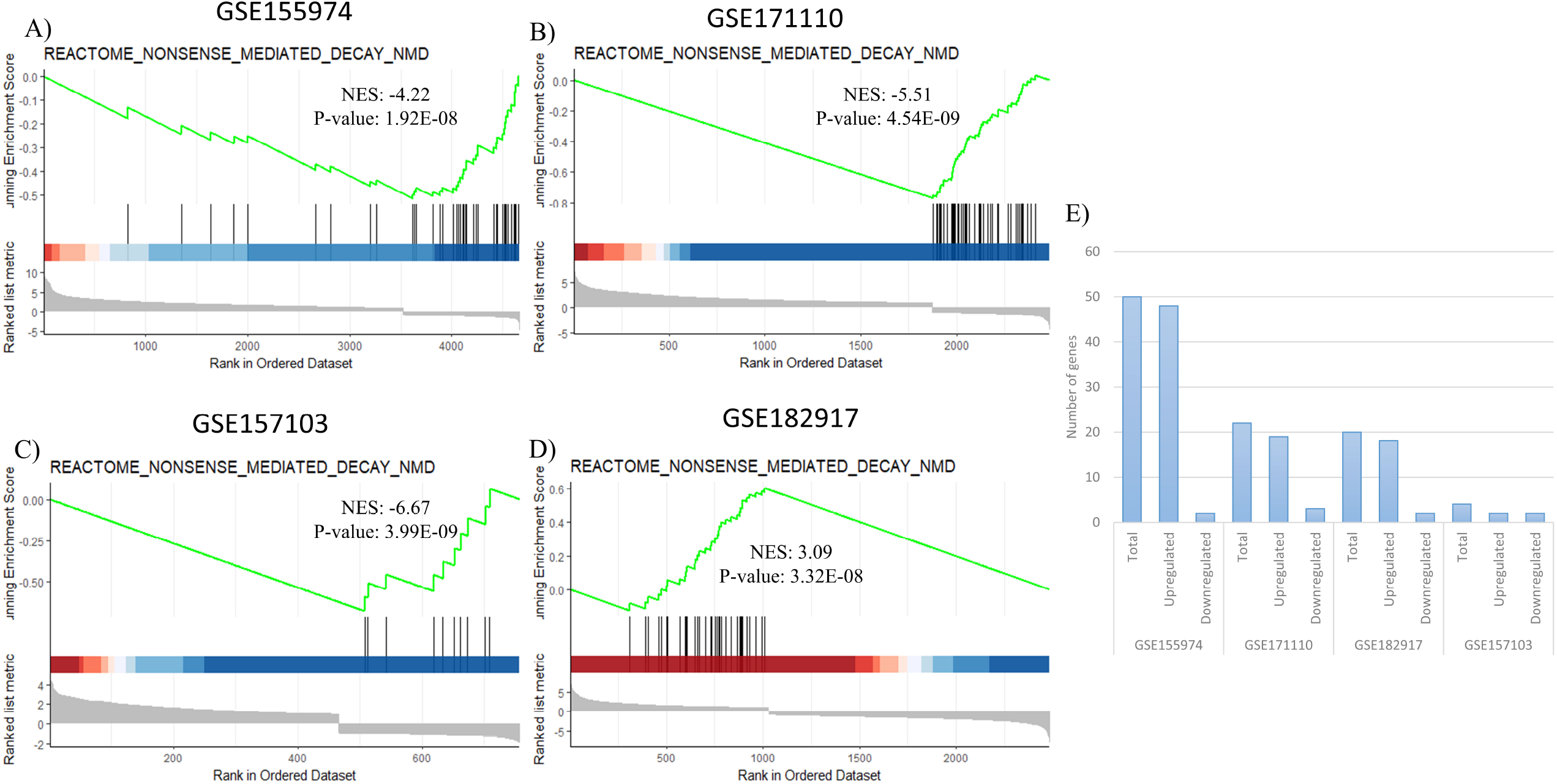
GSEA analyses on different SARS-CoV-2-infected cells/tissues. GSEA analyses of GSE155974 (A), GSE171110 (B), and GSE157103 (C) GSE182917 (D). All GSEA analyses, except one, show negative enrichment of NMD pathway in samples infected with SARS-CoV-2. Number of DEGs in each DEG list of GSEAs that matched with upregulated genes in consequence of UPF1 knockdown (E).

### SARS-CoV2-induced transcriptional changes show a significant overlap with transcriptional changes induced by UPF1 knockdown

Perturbed activity of NMD pathway leads to increased expression of its target transcripts. While, PTC-containing transcripts represent an important group of NMD targets, studies have shown that numerous normal transcripts are also recognized and targeted by NMD (12). Presence of features like uORFs, long 3’UTRs and some other non-PTC elements can make the transcripts susceptible to NMD degradation. In an effort to systematically identify NMD target transcripts, Colombo et al performed transcript profiling on cells in which different NMD players were knocked down (19). Their results showed that the majority of NMD’s ‘normal’ targets were protein-coding genes, followed by pseudogenes and non-coding RNAs. Considering the availability of these data, we asked whether transcriptional changes caused by SARS-COV-2 infection, had any overlap with transcriptional alterations observed by Colombo et al following the knockdown of NMD elements. To this end, we compared the list upregulated and down-regulated DEGs in Colombo et al dataset with SARS-COV-2-infected cells/tissues. As shown in Figure 5, a significant overlap was seen for Up-DEGS between Colombo et al UPF1 KD study and GSE155974 as well as GSE171110. While this is indirect evidence, it likely reflects alterations induced by SARS-COV-2 in NMD pathway activity.

**Figure 5.**
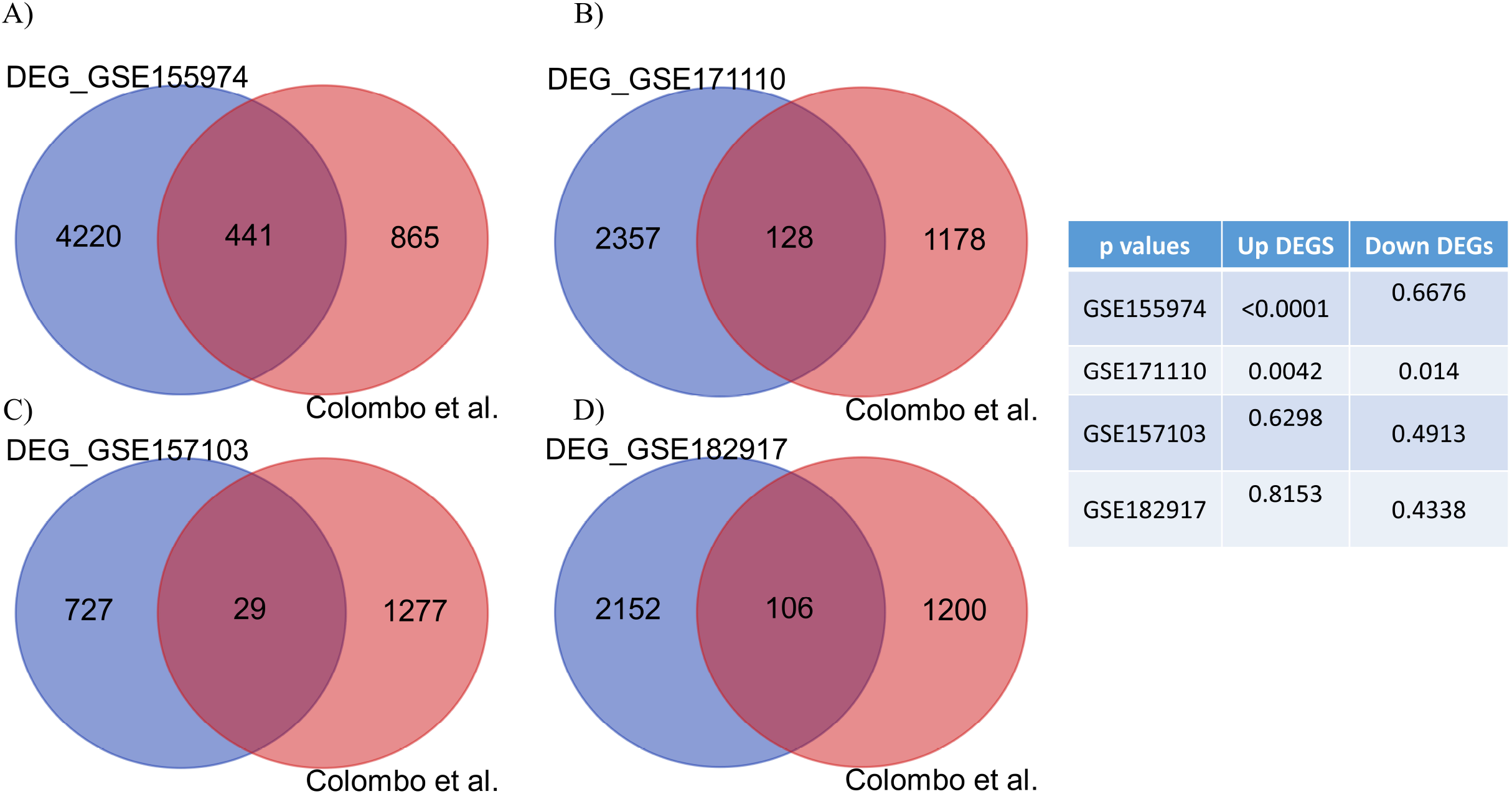
Venn diagrams showing the similarity in DEGs between Colombo et al study and SARS-CoV-2-infected cells/tissues. Transcripts showing significant alterations (FDR < 0.05) between Colombo et al study and GSE155974 (A), GSE171110 (B), and GSE157103 (C) GSE182917 (D) studies are shown. Table shows p values for Up and down-DEGs shared by the two studies.

## Discussion

Nonsense-mediated RNA decay (NMD) is known as an mRNA surveillance pathway that controls gene expression quality by recognizing and removing ‘faulty’ transcripts. NMD was initially discovered as an mRNA degradation pathway that detected transcripts with premature termination codons (PTCs) (20-22). Further studies revealed that NMD targets might have features other than PTCs; i.e. presence of long 3’ untranslated regions (UTR), upstream open reading frame (uORF) or termination codons which ‘mimic’ PTCs. NMD initiation requires a set of evolutionarily conserved proteins including up-frameshift protein 1 to 3 (UPF1-3) as well as SMG1/SMG5-9 which are involved in RNA cleavage and degradation (23, 24) and polypeptide chain release factors also known as eRFs (11, 25-27).

UPF1 is a helicase that is known as the master regulator of NMD in cells (28). It is also involved in other processes including DNA repair and replication, telomerase hemostasis, mRNA transport and RNA localization. This protein consists of three main regions; an N-terminal cysteine-histidine rich (CH) domain which binds to UPF2, ribosomal protein S26 (RPS26), and mRNA-decapping enzyme 2 (DCP2), a helicase domain that binds to single-stranded RNA (ssRNA) and DNA (ssDNA) molecules and a nonstructural serine-glutamine (SQ)-rich C-terminal domain which interact with SMGs (suppressor with morphogenetic defects in genitalia) (10, 29-32).

During translation of ‘normal’ transcripts, eukaryotic release factor 1 (eRF1) is required for recognition of stop codons and termination of translation. Following the recognition of stop codon by eRF1, another protein named eRF3 binds to eRF1 and facilitates the release of the nascent polypeptide from the ribosome through its GTPase activity (33). Another player is cytoplasmic polyA-binding protein 1 (PABPC1) which binds to mRNA polyA tails and enhances the recruitment of eRFs (34). During translation, movement of ribosomes across the transcripts leads to the displacement and removal of exon junction complex (EJC) proteins which are normally present at EJCs. In an aberrant mRNA translation termination; e.g. when the ribosome reaches a termination codon which is located upstream of an EJC, the presence of unremoved EJC proteins leads to the recruitment of UPF proteins to the location. UPF1 is recruited via UPF2 to EJC-UPF3B complex and is phosphorylated by SMGs, which finally results in NMD pathway execution (27, 35). NMD activation might also happen in the absence of EJC proteins. In one of the mechanisms reported for EJC-independent NMD activation, the transcript’s long 3’-UTR can prevent PABPC1 from interacting with eRF3 and instead allowing UPF1 interaction to eRF3 (36). Hence, the competition between PAPBC1 and UPF1 for binding to eRF3 can change the mRNA survival (35, 37).

Many viruses use long RNA transcripts containing multiple ORFs during their gene expression and life cycle. These long RNAs generally contain several termination codons inside the transcript and far from the polyA tail. This feature mimics the presence of PTCs in eukaryotic transcripts and can lead to the activation of NMD pathway and degradation of viral transcripts. That said, it has been shown that viruses employ different mechanisms to protect their transcripts from the NMD pathway (27). Semliki Forest virus (SFV) and Sindbis virus (SINV) are alphaviruses from Togaviridae family which carry positive-sense ssRNA with long 3’-UTRs. Genome-wide small interfering RNA (siRNA) studies used to identify host factors involved in viral replication have shown that UPF1 and other NMD factors act as restrictors for the replication of these viruses by degrading their RNAs (38, 39). A combination of proteomics and RNAi screening approaches on hepatitis C virus (HCV) infected cells have also revealed that the core protein of HCV inhibits NMD and increases virus replication in hepatoma cell lines (40). NMD-inhibiting mechanisms have also been reported for retroviruses. Rous Sarcoma Virus (RSV) has a 400 nucleotide long sequence downstream of its gag mRNA called RNA stability element (RSE) which can form an RNA secondary structure and prevent the recruitment of UPF1 and the execution of NMD (41, 42). Moreover, tax and rex, two proteins from Human T-lymphotropic Virus Type 1 (HTLV-1) have been shown to bind UPF1 and prolong viral genome half-life by preventing their RNA from degradation (43, 44). A few studies have demonstrated interactions between HIV-1 with UPF proteins (45-47). There is evidence that anti-NMD mechanisms might also exist for coronaviruses. In a study by Wada et al researchers have shown that N protein of murine hepatitis virus (MHV), a member of the betacoronaviruses, may partially inhibit NMD execution (48). Whether other MHV or coronaviral proteins can inhibit NMD is unclear.

In this study, we found significant sequence and structure similarity between SARS-CoV-2 helicase (nsp13 in other pathogenic β-CoVs) with human UPF1. We then used GSEA analysis of RNA-seq data derived from SARS-Cov2-infected samples to see whether NMD-related genes show any alterations following viral infection. We then examined transcriptomic data from UPF1 knockout cells to investigate whether any similarities might exist between these cells and virus infected samples at the molecular level. Our analyses provided some evidence, albeit indirect, for possible interference of viral helicase (nsp13) with the NMD pathway. It is conceivable that SARS-CoV-2 helicase could interfere with cellular NMD by interacting with UPF2 and eRF through its sequence and structural similarity with UPF1. If supported by experimental evidence, this might indicate the presence of a virus-host interaction that compromises an important cell-intrinsic defense mechanism.

## Acknowledgments

Not applicable.

## Supporting information

**Table S1**. SARS-CoV-2 protein’s accession number and symbol.

**Table S2**. Delta-BLAST parameters.

**Table S3**. GEO Dataset attributes.

**Figure S1. Protein-protein interaction map for UPF1**. UPF1 interacts with CASC3, UPF2, GSPT2, UPF3A, EIF4A3, GSPT1, UPF3B, SMG7, RBM8A, SMG1 in Homo Sapiens (A). Gene Ontology Map showing important biological processes (BP), cellular components (CC), and molecular functions (MF) in cells (B). GO analysis of UPF1 and its interacting proteins revealed these proteins main biological process in cells is non-sense RNA mediated decay.

## References

1. Cui J, Li F, Shi ZL. Origin and evolution of pathogenic coronaviruses. Nat Rev Microbiol. 2019;17(3):181–92.

2. Andersen KG, Rambaut A, Lipkin WI, Holmes EC, Garry RF. The proximal origin of SARS-CoV-2. Nat Med. 2020;26(4):450–2.

3. Davey NE, Trave G, Gibson TJ. How viruses hijack cell regulation. Trends Biochem Sci. 2011;36(3):159–69.

4. Tarakhovsky A, Prinjha RK. Drawing on disorder: How viruses use histone mimicry to their advantage. J Exp Med. 2018;215(7):1777–87.

5. Alhammad YMO, Fehr AR. The Viral Macrodomain Counters Host Antiviral ADP-Ribosylation. Viruses. 2020;12(4).

6. Fairman-Williams ME, Guenther UP, Jankowsky E. SF1 and SF2 helicases: family matters. Curr Opin Struct Biol. 2010;20(3):313–24.

7. May JP, Simon AE. Targeting of viral RNAs by Upf1-mediated RNA decay pathways. Curr Opin Virol. 2021;47:1–8.

8. Han W, Li X, Fu X. The macro domain protein family: structure, functions, and their potential therapeutic implications. Mutat Res. 2011;727(3):86–103.

9. Holm L. Using Dali for Protein Structure Comparison. Methods Mol Biol. 2020;2112:29–42.

10. Gowravaram M, Bonneau F, Kanaan J, Maciej VD, Fiorini F, Raj S, et al. A conserved structural element in the RNA helicase UPF1 regulates its catalytic activity in an isoform-specific manner. Nucleic Acids Res. 2018;46(5):2648–59.

11. Lykke-Andersen S, Jensen TH. Nonsense-mediated mRNA decay: an intricate machinery that shapes transcriptomes. Nat Rev Mol Cell Biol. 2015;16(11):665–77.

12. Hug N, Longman D, Caceres JF. Mechanism and regulation of the nonsense-mediated decay pathway. Nucleic Acids Res. 2016;44(4):1483–95.

13. Nasif S, Contu L, Muhlemann O. Beyond quality control: The role of nonsense-mediated mRNA decay (NMD) in regulating gene expression. Semin Cell Dev Biol. 2018;75:78–87.

14. Leon K, Ott M. An ‘Arms Race’ between the Nonsense-mediated mRNA Decay Pathway and Viral Infections. Semin Cell Dev Biol. 2021;111:101–7.

15. Popp MW, Cho H, Maquat LE. Viral subversion of nonsense-mediated mRNA decay. RNA. 2020;26(11):1509–18.

16. Knoll R, Schultze JL, Schulte-Schrepping J. Monocytes and Macrophages in COVID-19. Front Immunol. 2021;12:720109.

17. Percivalle E, Sammartino JC, Cassaniti I, Arbustini E, Urtis M, Smirnova A, et al. Macrophages and Monocytes: “Trojan Horses” in COVID-19. Viruses. 2021;13(11).

18. Balistreri G, Horvath P, Schweingruber C, Zund D, McInerney G, Merits A, et al. The host nonsense-mediated mRNA decay pathway restricts Mammalian RNA virus replication. Cell Host Microbe. 2014;16(3):403–11.

19. Colombo M, Karousis ED, Bourquin J, Bruggmann R, Muhlemann O. Transcriptome-wide identification of NMD-targeted human mRNAs reveals extensive redundancy between SMG6- and SMG7-mediated degradation pathways. RNA. 2017;23(2):189–201.

20. Kurosaki T, Maquat LE. Nonsense-mediated mRNA decay in humans at a glance. Journal of Cell Science. 2016;129(3):461–7.

21. Brogna S, Wen J. Nonsense-mediated mRNA decay (NMD) mechanisms. Nat Struct Mol Biol. 2009;16(2):107–13.

22. Hug N, Longman D, Cáceres JF. Mechanism and regulation of the nonsense-mediated decay pathway. Nucleic Acids Res. 2016;44(4):1483–95.

23. Lykke-Andersen S, Chen Y, Ardal BR, Lilje B, Waage J, Sandelin A, et al. Human nonsense-mediated RNA decay initiates widely by endonucleolysis and targets snoRNA host genes. Genes Dev. 2014;28(22):2498–517.

24. Boehm V, Haberman N, Ottens F, Ule J, Gehring NH. 3’ UTR length and messenger ribonucleoprotein composition determine endocleavage efficiencies at termination codons. Cell Rep. 2014;9(2):555–68.

25. Rebbapragada I, Lykke-Andersen J. Execution of nonsense-mediated mRNA decay: what defines a substrate? Curr Opin Cell Biol. 2009;21(3):394–402.

26. Schweingruber C, Rufener SC, Zünd D, Yamashita A, Mühlemann O. Nonsense-mediated mRNA decay - mechanisms of substrate mRNA recognition and degradation in mammalian cells. Biochim Biophys Acta. 2013;1829(6-7):\p612-23.

27. Balistreri G, Bognanni C, Mühlemann O. Virus Escape and Manipulation of Cellular Nonsense-Mediated mRNA Decay. Viruses. 2017;9(1).

28. Fiorini F, Bagchi D, Le Hir H, Croquette V. Human Upf1 is a highly processive RNA helicase and translocase with RNP remodelling activities. Nat Commun. 2015;6:7581.

29. He F, Brown AH, Jacobson A. Upf1p, Nmd2p, and Upf3p are interacting components of the yeast nonsense-mediated mRNA decay pathway. Mol Cell Biol. 1997;17(3):1580–94.

30. Min EE, Roy B, Amrani N, He F, Jacobson A. Yeast Upf1 CH domain interacts with Rps26 of the 40S ribosomal subunit. Rna. 2013;19(8):1105–15.

31. He F, Jacobson A. Identification of a novel component of the nonsense-mediated mRNA decay pathway by use of an interacting protein screen. Genes Dev. 1995;9(4):437–54.

32. Denning G, Jamieson L, Maquat LE, Thompson EA, Fields AP. Cloning of a novel phosphatidylinositol kinase-related kinase: characterization of the human SMG-1 RNA surveillance protein. J Biol Chem. 2001;276(25):22709–14.

33. Nakamura Y, Ito K, Matsumura K, Kawazu Y, Ebihara K. Regulation of translation termination: conserved structural motifs in bacterial and eukaryotic polypeptide release factors. Biochem Cell Biol. 1995;73(11-12):1113–22.

34. Ivanov A, Mikhailova T, Eliseev B, Yeramala L, Sokolova E, Susorov D, et al. PABP enhances release factor recruitment and stop codon recognition during translation termination. Nucleic Acids Res. 2016;44(16):7766–76.

35. Singh G, Rebbapragada I, Lykke-Andersen J. A competition between stimulators and antagonists of Upf complex recruitment governs human nonsense-mediated mRNA decay. PLoS Biol. 2008;6(4):e111.

36. Bhuvanagiri M, Schlitter AM, Hentze MW, Kulozik AE. NMD: RNA biology meets human genetic medicine. Biochem J. 2010;430(3):365–77.

37. Mühlemann O, Jensen TH. mRNP quality control goes regulatory. Trends Genet. 2012;28(2):70–7.

38. Moon SL, Wilusz J. Cytoplasmic viruses: rage against the (cellular RNA decay) machine. PLoS Pathog. 2013;9(12):e1003762.

39. Balistreri G, Horvath P, Schweingruber C, Zünd D, McInerney G, Merits A, et al. The host nonsense-mediated mRNA decay pathway restricts Mammalian RNA virus replication. Cell Host Microbe. 2014;16(3):403–11.

40. Ramage HR, Kumar GR, Verschueren E, Johnson JR, Von Dollen J, Johnson T, et al. A combined proteomics/genomics approach links hepatitis C virus infection with nonsense-mediated mRNA decay. Mol Cell. 2015;57(2):329–40.

41. Withers JB, Beemon KL. The structure and function of the rous sarcoma virus RNA stability element. J Cell Biochem. 2011;112(11):3085–92.

42. Ge Z, Quek BL, Beemon KL, Hogg JR. Polypyrimidine tract binding protein 1 protects mRNAs from recognition by the nonsense-mediated mRNA decay pathway. Elife. 2016;5.

43. Mocquet V, Neusiedler J, Rende F, Cluet D, Robin JP, Terme JM, et al. The human T-lymphotropic virus type 1 tax protein inhibits nonsense-mediated mRNA decay by interacting with INT6/EIF3E and UPF1. J Virol. 2012;86(14):7530–43.

44. Nakano K, Ando T, Yamagishi M, Yokoyama K, Ishida T, Ohsugi T, et al. Viral interference with host mRNA surveillance, the nonsense-mediated mRNA decay (NMD) pathway, through a new function of HTLV-1 Rex: implications for retroviral replication. Microbes Infect. 2013;15(6-7):491–505.

45. Ajamian L, Abrahamyan L, Milev M, Ivanov PV, Kulozik AE, Gehring NH, et al. Unexpected roles for UPF1 in HIV-1 RNA metabolism and translation. Rna. 2008;14(5):914–27.

46. Ajamian L, Abel K, Rao S, Vyboh K, García-de-Gracia F, Soto-Rifo R, et al. HIV-1 Recruits UPF1 but Excludes UPF2 to Promote Nucleocytoplasmic Export of the Genomic RNA. Biomolecules. 2015;5(4):2808–39.

47. Serquiña AK, Das SR, Popova E, Ojelabi OA, Roy CK, Göttlinger HG. UPF1 is crucial for the infectivity of human immunodeficiency virus type 1 progeny virions. J Virol. 2013;87(16):8853–61.

48. Wada M, Lokugamage KG, Nakagawa K, Narayanan K, Makino S. Interplay between coronavirus, a cytoplasmic RNA virus, and nonsense-mediated mRNA decay pathway. Proc Natl Acad Sci U S A. 2018;115(43):E10157–e66.

