## Supplementary figures and images for "SARS-CoV-2 Helicase might interfere with cellular nonsense-mediated RNA decay, insights from a bioinformatics study"

### Supplemental Figure 1

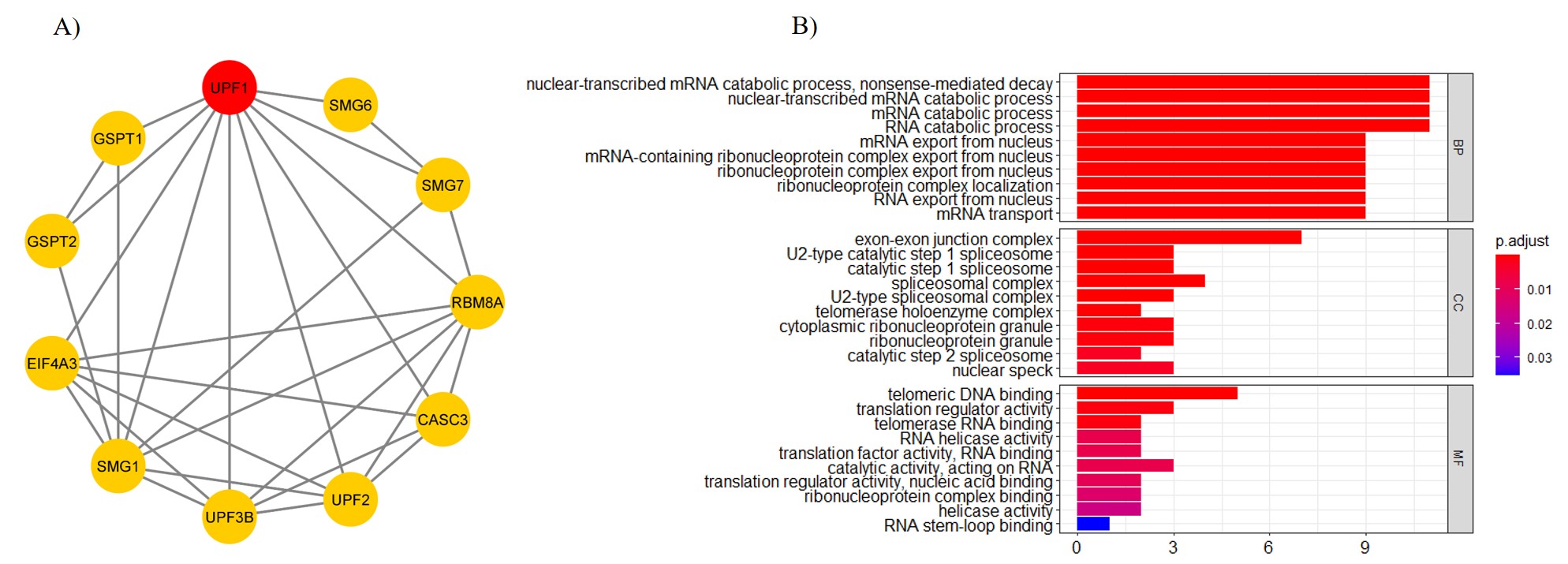

### Supplemental Table 1

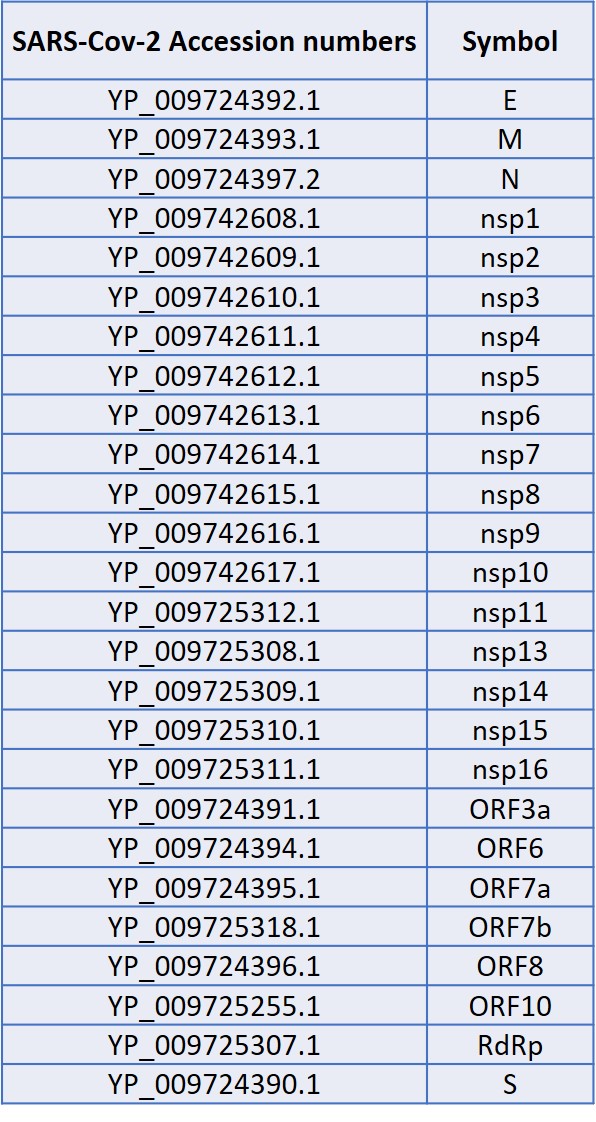

### Supplemental Table 2

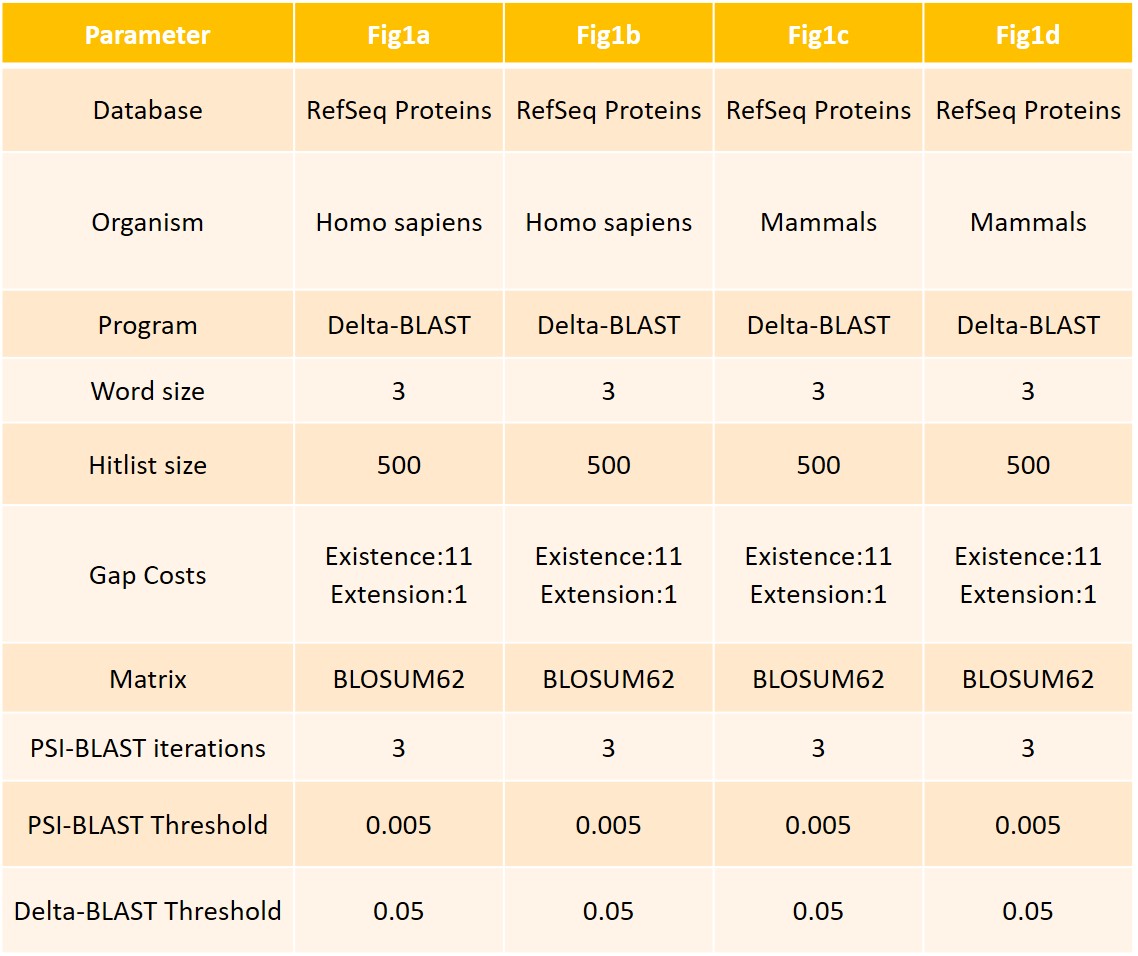

### Supplemental Table 3

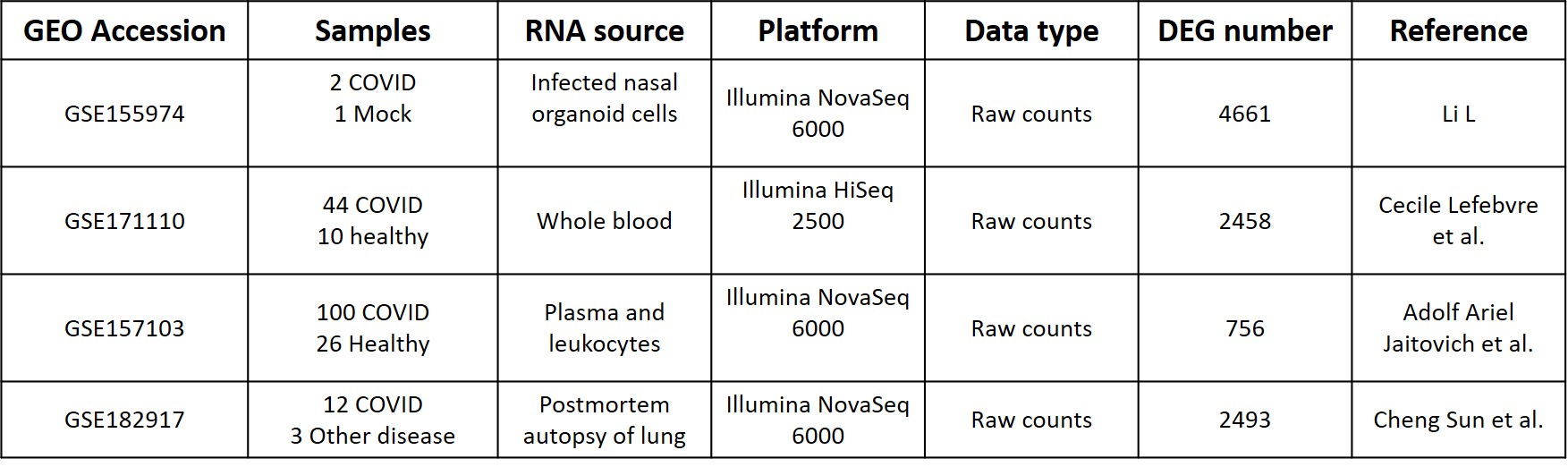
